# Label-free super-resolution chemical imaging of biomedical specimens

**DOI:** 10.1101/2021.05.14.444185

**Authors:** Julien Guilbert, Awoke Negash, Simon Labouesse, Sylvain Gigan, Anne Sentenac, Hilton B. de Aguiar

**Affiliations:** Laboratoire Kastler Brossel, ENS-Université PSL, CNRS, Sorbonne Université, Collège de France, 24 rue Lhomond, 75005 Paris, France; Aix Marseille Univ, CNRS, Centrale Marseille, Institut Fresnel, Marseille, France

## Abstract

Raman microscopy provides chemically selective imaging by exploiting intrinsic vibrational properties of specimens. Yet, a fast acquisition, low phototoxicity, and non-specific (to a vibrational/electronic mode) super-resolution method has been elusive for tissue imaging. We demonstrate a single-pixel-based approach, combined with robust structured illumination, that enables fast super-resolution in stimulated Raman scattering microscopy at low power levels. The methodology is straightforward to implement and compatible with thick biological specimens, therefore paving the way for probing complex biological systems when exogenous labelling is challenging.

---

Far-field super-resolution imaging has emerged as a powerful tool in cell biology to unravel the details of the complex molecular machinery at play at the nanoscale. However, the great majority of super-resolution techniques are based on exogenous markers (fluorophores) that demand careful preparation protocols to determine cell viability and specificity to a targeted molecule. Most importantly, fluorescence-based tools only report on the fluorophore information – dynamical or structural – leaving open many fundamental questions on the other out- numbering unlabelled molecular species: e.g., lipids and cholesterol molecular conformation and local composition [1, 2] within lipid rafts domains have remained undetected in real cells, or the local composition of the species forming membrane-less organelles which are currently unknown. Therefore, Raman microscopies have emerged as ideal tools for probing heterogeneous biological specimens [3], since they provide chemically resolved images using the intrinsic vibrational properties of molecules. Yet, reaching fast super-resolution capabilities in Raman microscopies has remained elusive [4].

In the last decade, many attempts have been made to enable vibrational far-field super-resolution. To date, computational super-resolution methods, exploiting structured illumination microscopy (SIM), have been demonstrated for the spontaneous Raman case [5]. However, the usage of an imaging spectrometer is not compatible with thick tissues as the resolution enhancement is only provided in one dimension, and the acquisition speeds are too low for dynamic specimens. Alternatively, coherent Raman microscopies (CRM) — with Coherent anti-Stokes Raman scattering (CARS) [6] and stimulated Raman scattering (SRS) [7–9] being the two most known contrast mechanisms in CRM — could provide fast acquisition speeds, however there are various drawbacks that preclude biological specimens imaging. For CARS, interference artifacts complicate chemical quantification analysis [10]. In the case of the quasi background-free SRS process, the current mainstream is to exploit methods to control the dynamics of vibrational energy levels, however using unconventional power levels that may be phototoxic for biological specimens [11].

Ideally, combining computational super-resolution methods with SRS technology would overcome the above-mentioned issues, but is currently elusive. Generally, the mathematical framework of computational methods are based on wide-field geometries having no assumptions on the power levels needed, yet requiring multipixel cameras. Unfortunately, wide-field cameras for SRS are technologically challenging because of SRS’ current paradigm detection scheme involving single-pixel detectors with high-sensitivity radio-frequency lock-in amplifier (RF-LIA). Despite recent developments of multi-pixel RF-LIA [12], the pixel counts do not scale favorably for 1000’s of pixels architecture needed in a camera. Furthermore, on a more fundamental note, wide-field geometries are not suitable for thick tissue imaging due to the lack of sectioning capabilities.

Here we present a chemically selective imaging methodology compatible with thick biological specimens that breaks the diffraction-limit resolution barrier of SRS microscopy. Specifically, we developed a single-pixel method (Fig. 1) compatible with computational super-resolution methods, therefore allowing for fast imaging capabilities exploiting SRS processes in the form of stimulated Raman gain (SRG) (Fig. 1a). In our scheme, a structured stationary pump beam is shaped using a spatial light modulator (SLM) and is spatially and temporally overlapped with a focused Stokes beam that is scanned over (using a set of galvanometric mirrors, Fig. 1b). This scanning scheme generates an image that is of lower resolution than an image performed by a conventional SRS microscope (Fig. 1c). After acquiring a series of SRS images with multiple structured illuminations the data is treated with algorithms based on standard SIM mathematical framework to recover a super-resolved image [13, 14] (see Supplementary Materials SI for a complete discussion of the image formation in our methodology). Remarkably, the framework presented here has a simple alignment procedure: it is simpler than conventional SRS microscopy, which demands overlap of two tightly focused beams. Furthermore, we chose to work with non-sinusoidal SIM patterns in order to be compatible with thick tissues: we use speckle patterns, since they are resilient in scattering specimens. We coin the method Single-pixel blind-SIM SRS (or blind-S^3^ for short).

**FIG. 1.**
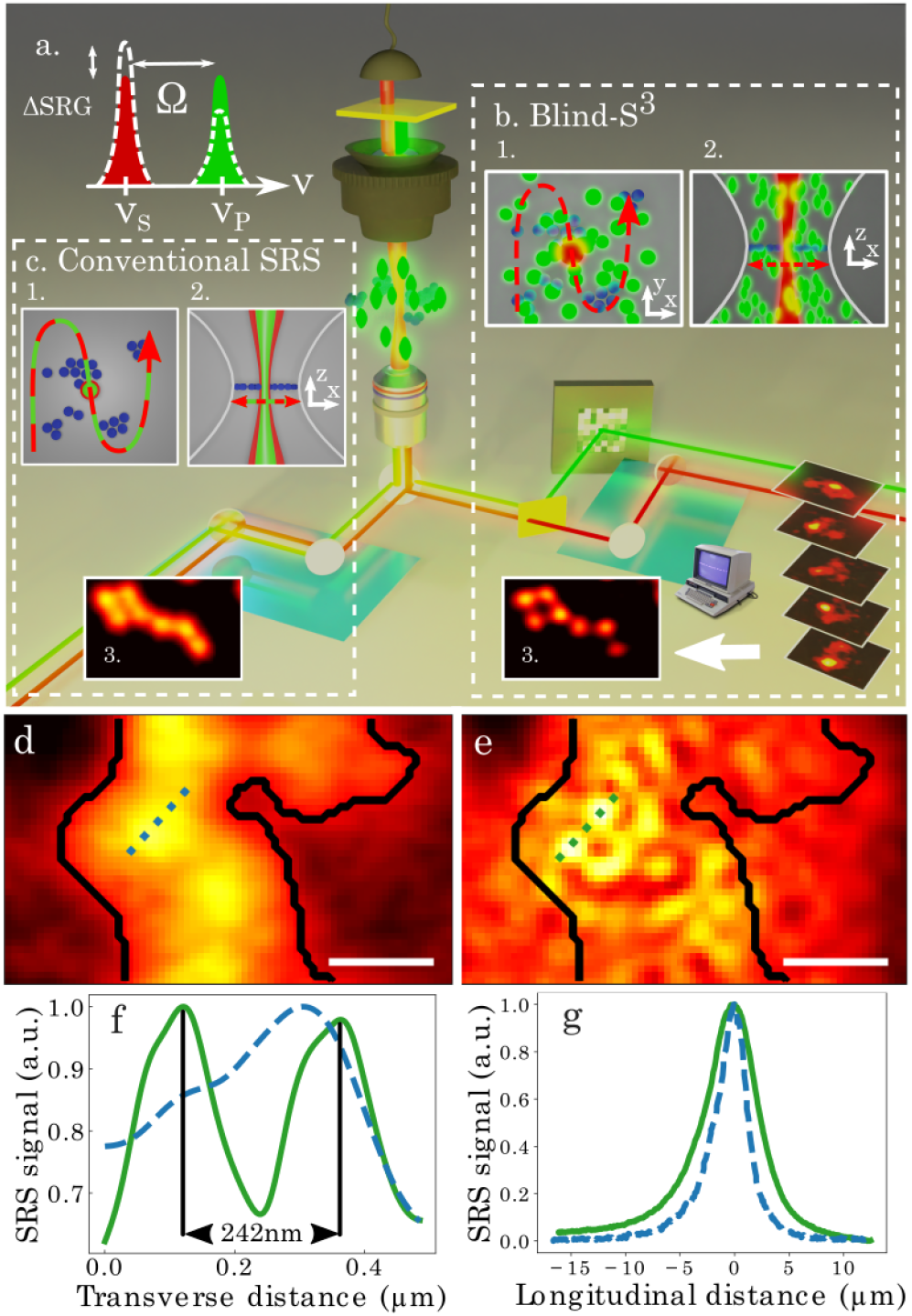
Principle of blind-S3 and demonstration of imaging beyond the conventional SRG diffraction-limit. Schematic of the setup to achieve super resolution using SRG process (**a**) based on single-pixel SIM scheme. Transverse (**b**1) and longitudinal (**b**2) planes of the Stokes beam (red dash) scanning trajectory over the stationary Raman- active molecules (blue) and structured Pump (green), in this case a speckle pattern. For every speckle realization, an SRG image is acquired forming a stack that is passed to a SIM algorithm to reconstruct a super-resolved image (**b**3). (**c**) Conventional SRG, consisting in raster scanning co-propagating Pump and Stokes beams, is used as a control to demonstrate the increase in resolution when compared to standard imaging. Transverse (**c**1) and longitudinal (**c**2) planes of the Stokes beam and focused Pump (green and red dash) scanning trajectory over the stationary Raman-active molecules (blue). Conventional (**d**) and blind-S^3^ (**e**) images of 239 nm-diameter polystyrene beads, and line profiles (**f**) showing the increase in transverse resolution SRG (dash) and blind-S^3^ (line). (**g**) Conventional SRG (dash) and blind-S^3^ (line) sectioning capabilities characterization. All scale bars: 500 nm.

We first demonstrate the improvement in the transverse resolution, surpassing the usual diffraction-limit of SRS microscopy. In order to evaluate the gain in resolution, we compare blind-S^3^ to the conventional scanning methods. For conventional SRS, the theoretical transverse resolution is Δ*r*^*Conv*^ = 307 nm (see Methods for more detail). This theoretical value is technically challenging to achieve with high NA objectives in the near-IR because the wavelengths of the two beams differ by hundreds of nanometers (spectral span necessary for fast quantification of lipids, proteins, and nucleic acids in SRS microscopy). Conversely, blind-S^3^ transverse spatial resolution results from the doubling in resolution dictated by SIM and the speckle grain size limited by diffraction, leading to 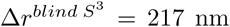 (see Methods for more detail). To show the superior transverse resolution in blind-S^3^, we imaged 239 nm-diameter polystyrene beads with the two modalities (Fig. 1d-e). Clearly, conventional SRS (Fig. 1d) cannot resolve the beads transversely as the beads size is smaller than the theoretical resolution limit. After multiple speckle pattern illuminations, we feed the resulting images to a blind SIM algorithm to reconstruct a super-resolved image (see Methods for details). Remarkably, blind-S^3^ methodology (Fig. 1e) resolves several beads in the in-focus layer. The line profiles reveal the distance between the centers of the beads (242 nm) which matches well to the distance of close contact between two beads. We note that the effective ROI of blind-S^3^ is modulated by the speckle envelope, hence decreasing the similarity of the two images in the edges of the beads cluster, yet not affecting the resolution gain (a resolution quantification of multiple beads size and NA is presented in Supplementary Materials SII). The present findings show that the blind-S^3^ methodology goes beyond the fundamental far-field diffraction-limit resolut<u>i</u>on of SRS microscopy, by improving the resolution 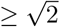.

Remarkably, and contrary to conventional wide-field imaging, super-resolution in blind-S^3^ comes for free with high z-sectioning capabilities. In each illumination during the blind-S^3^ procedure, an image is formed based on a wide-field geometry model, that is, an object is convoluted with a linear point-spread function (PSF). In the conventional wide-field microscope using multi-pixel cameras, the acquisition procedure is not able to effectively reject signal from out-of-focus region, therefore deteriorating image quality and resolution due to the background shot-noise. Conversely, in blind-S^3^ the nonlinear optical response is local in the longitudinal direction, because SRS signal is only generated within the overlap region of the two beams, hence each SRS image illumination does not contain appreciable out-of-focus signal. To demonstrate the sectioning capabilities of blind-S^3^, despite an image being a convolution with a linear PSF, we probed a thin film of oil (of few *µ*m) by scanning it in the longitudinal direction. Note that we collect the signal generated for each z-position on a *≈* 10 mm-wide detector, hence, not in a confocal geometry. Clearly, conventional SRS and blind-S^3^ give a peaked response which means that those two techniques have inherent longitudinal sectioning (Fig. 1g). Indeed, the conventional SRS microscope is able to show such z-sectioning capability, due to its nonlinear longitudinal PSF. Different from conventional SRS, in blind-S^3^ the z-sectioning is coupled with the effective field-of-view (FOV), with a weak dependence, since the speckle envelope in the transverse direction is coupled with the longitudinal one (Supplementary Materials SIII presents detailed explanation of the sectioning in blind-S^3^). We further show in the Supplementary Materials SIV that the resolution in the z-direction for both techniques are similar.

To demonstrate blind-S^3^ compatibility with biological specimens, we image standard cell lines and mouse brain tissues. Conventional SRS reveals several *µ*m-large droplets within the cell in the FOV (Fig. 2a). Close-up images show different cluster morphology (Fig. 2b), and increased resolution gain with blind-S^3^ from the line profiles of selected ROI (Fig. 2c). To demonstrate capabilities for aberrant and opaque tissues, we have further imaged highly scattering brain slices at 8 *µ*m-deep in the sample (Fig. 2d-e) with line profiles demonstrating increased resolution power of the myelin structures (Fig. 2f). The close-up images with super-resolution capabilities (Fig. 2e) reveal that the structure of the myelin in the tissue is actually not as symmetrically perfect as inferred from the low-resolution images. These results show that the method is fully compatible with thick tissue imaging, despite being opaque.

**FIG. 2.**
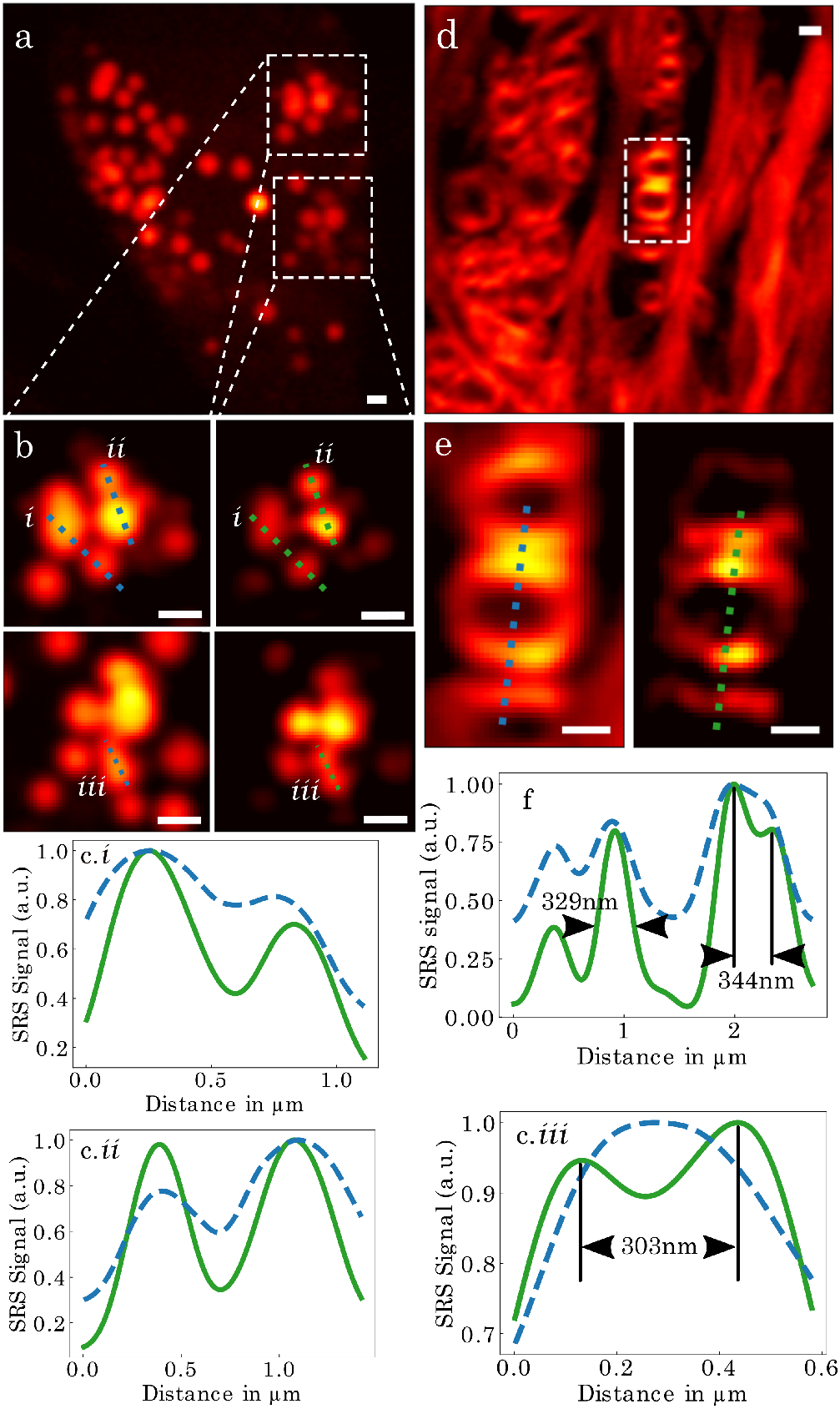
Bio-compatibility capabilities of blind-S^3^. (**a**) Large FOV imaging of lipid droplets within HeLa cells (conventional SRS). Two zoomed-in ROIs (**b**) are depicted by dashed boxes with conventional SRS (left panels) and blind- S^3^ (right panels) methods, with various line profiles shown (c, **i, ii** and **iii**) for conventional SRS (dash) and blind-S^3^ (line). (**d**) Large FOV image of 100 μm thick mouse cerebellum (conventional SRS). A zoomed-in ROI (**e**) is depicted by dashed boxes with conventional SRS (left panel) and blind-S^3^ (right panel) methods, with a line profile chosen (**f**) for conventional (dash) and blind-S^3^ (line). All scale bars: 500 nm.

We have designed and demonstrated a single-pixel super-resolution technique that is straightforward to implement, i.e. simpler than a conventional SRS microscope. Furthermore, our method is able to image opaque biological tissues, meaning that it would be compatible for epi imaging typical of *in-vivo* situations [3]. Blind-S^3^ is a universal approach in the sense that it does not depend on the specific vibrational mode ultrafast dynamics [11] and does not require *a priori* knowledge about the specimen, for instance, as gained in the training of ne<u>u</u>ral networks methods [15]. We demonstrate a gain of 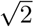 in transverse resolution with high z-sectioning capabilities. We anticipate that far-field nanoscopy (sub-100-nm resolution) may be achieved by using visible wavelengths, where high-NA objectives perform much better [16], without compromising signal level. blind-S^3^ not only paves the way for fast super-resolution in Raman imaging but open up a series of other developments in super-resolution microscopy of biological tissues, since the assumptions used are very similar to the ones found in multi-photon fluorescence microscopy.

## METHODS

### Microscope design and details

A detailed figure of the setup is presented in Supplementary Materials V. Briefly, the output power of a femtosecond laser source (Coherent, Chameleon Ultra Vision, 800 nm, 80 MHz repetition rate, 150 fs pulse length) pumps an optical parametric oscillator (APE, MIRA-OPO) that generates the Stokes beam, centered either at 1058 nm (Fig. 1, 3054 cm^*-*1^ Raman-shift) or 1048 nm (Fig. 2, 2958 cm^*-*1^ Raman-shift), and a small power fraction is used as the Pump beam. The Stokes beam is spectrally narrowed using a combination of grating (LightSmyth, T-1000-1040) and adjustable slit width for the purpose of increasing chemical selectivity. The Pump beam is also spectrally narrowed in a pulse-shaper setup using two gratings (LightSmyth Technologies, T-1400-800) and a digital micromirror device (DMD) placed in the Fourier plane (a description of the methods using DMDs for SRS spectroscopy can be found in Ref. [17]). The pump beam is amplitude-modulated at 1 MHz by an acousto-optic modulator (AA Opto-electronic, MT80-B30A1,5 VIS). The specimen is z-displaced using a piezo stage (Thorlabs, DRV517), and the signal generated by the sample is then collected by a 1.4 NA oil-immersion condenser, directed to a large-area detector (Thorlabs, DET100A2) and demodulated by a lock-in amplifier (Zurich Instruments, MFLI).

We used two configurations for SRS microscopy. Regardless of the configuration used, both beams are spatially and temporally combined at dichroic mirrors, whose location depends on the modality of SRS in use, and focused by an objective (Nikon, Plan APO IR, 60x, NA=1.27). To achieve the best compromise in terms of resolution enhancement (by having the Pump beam as a structured illumination) and sensitivity (by having the Stokes beam as the demodulated beam), we have designed a layout that allows us to quickly swap the direction of the Pump beam between the conventional SRS or blind-S^3^ configurations using a combination of a halfwave waveplate and a polarizing beam splitter cube. For the blind-S^3^ configuration, the Pump beam is sent onto a SLM (Meadowlark Optics, HSP512L-1300) to modulate the wavefront with a random phase, thus generating a speckle pattern at the image plane with user-selectable FOV. Galvanometric mirror scanners are used to move either Pump and Stokes beams together (conventional) or Stokes only (blind-S^3^). Typical average power measured before the objectives were 13 mW (conventional) and 41 mW (blind-S^3^) for the Pump, and 25 mW for the Stokes beams. However, we note that the energy density levels used for blind-S are inherently lower than the 2 conventional SRS configuration: we have estimated a*≈* 5 times lower effective energy densities (i.e. product of the energy densities of the Pump and Stokes energy densities), taking into consideration the speckle envelope and the longer integration time in the blind-S^3^ procedure.

### Sample preparation

Samples presented in Fig 2 were prepared by drop-casting the polystyrene beads on a coverslip and embedded in deuterated water to decrease the spectral congestion with the water vibrational response background. The various diameters (and standard deviation) used were: 239 nm (6 nm, PS Research Particles), 372 nm (10 nm, Polysciences, Inc.), 520 nm (16 nm, Thermo Scientific), 740 nm (22 nm, Thermo Scientific) and 990 nm (30 nm, Polysciences, Inc.). Brain slices, sectioned into 100 *µ*m horizontal slices in the sagital plan, were cut and stored in PBS containing 4% paraformaldehyde. Prior to experiments, the slices were placed between two cover slips with a 120-*µ*m-thick spacer. HeLa cells were incubated with 400 *µ*M oleic acid, washed, fixed with 4% paraformaldehyde, and stored at 4^*°*^C before imaging.

### Computational methods

We have simulated the methodology with various algorithms. An image acquired with blind-S^3^ scheme obeys a forward model *M*_*i*_ = (*O× I*_*Speckle*_) ⊛*I*_*Stokes*_, where *M*_*i*_ is an SRS image from a single speckle realization, *O* is the optical response of the excited object (more precisely, *ℑ*{*χ*^(3)^}), *I*_*Speckle*_ is the spatial distribution of the speckle intensity at the Pump wavelength, *I*_*Stokes*_ is the effective PSF of the image formation system, and ⊛denotes a convolution operation. While we tested the methodology with two SIM algorithms using no prior knowledge on the structured patterns *I*_*Speckle*_ [13, 14], for the results presented we used the one described in Ref. [14].

### Resolution estimation

We assume that the theoretical transverse resolution results from the product of two focused Gaussian beams with two different wavelengths *λ*_*p*_ and *λ*_*s*_ for the Pump and Stokes wavelength respectively. Here we use the Raman resonance 3054 cm^*-*1^ and the Rayleigh criteria to asses the resolution limit of each beam: 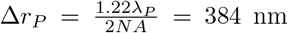 and 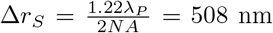 for the Pump and Stokes beam respectively where *NA* = 1.27, *λ*_*P*_ = 800 nm and *λ*_*S*_ = 1058 nm. Therefore, the theoretical resolution limit is 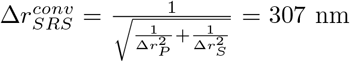 for conventional SRS while it is 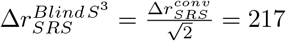 for blind-S^3^.

## Acknowledgements

We thank Claudio Moretti, Nabil Garroum, Arnaud Leclercq and Quentin Anty for technical support on the microscope development, Bernhard Rauer for fruitful discussions, Abdou Rachid Thiam and Mohyeddine Omrane for providing the HeLa cell lines, and Laurent Bourdieu for providing the brain slices. This research has been funded by the LabEX ENS-ICFP (ANR-10-LABX-0010/ANR-10-IDEX-0001-02 PSL*), FET-Open (Dynamic - 863203), European Research Council ERC Consolidator Grant (SMARTIES - 724473). S.G. is a member of the Institut Universitaire de France.

## Author contributions

All authors discussed the results.

## Competing interests

The authors declare no competing financial interests.

## Supplementary information

### I. ANALYTICAL FORWARD MODEL FOR BLIND-S^3^

In SRS, the signal detected (Δ*I*_*S*_) at one pixel location is given by:

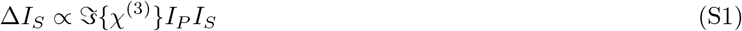

where *ℑ*{*χ*^(3)^} is the imaginary part of the complex-valued nonlinear susceptibility of the sample (related to the Raman cross-section), *I*_*P*_ and *I*_*S*_ the intensity of the Pump and Stokes beam, respectively.

In the case of blind-S^3^, a static speckle pattern generated by the Pump beam spreads at the sample image plane where the Stokes beam is focused and scanned. To derive an image formation model, we assume a scalar approximation for the local intensity in one blind-S^3^ image:

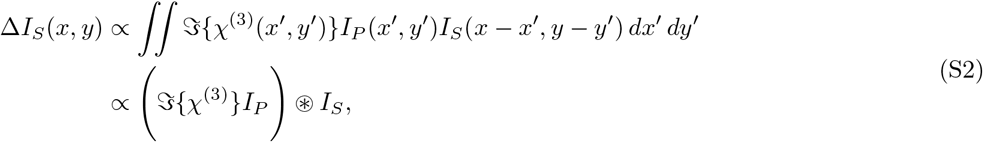

where ⊛ denotes the convolution operator, (*x, y*) the pixel position in the image Δ*I*_*S*_. In this approximation, we disregard coherent effects as SRS processes are inherently phase-matched.

This so-called forward model presented in Eq. S2 is mathematically similar to commonly used models in blind SIM reconstruction algorithms [S13, S14].

### II. TRANSVERSE RESOLUTION ANALYSIS

Contrary to conventional methods in super-resolution microscopy, in blind-S^3^ it is not straightforward to compare the reconstructed images with a “ground truth” object. This arises from the fact that the FOV in blind-S^3^ is determined by the speckle envelope, which is much smaller than conventional SRS microscopy. Therefore, we devised another methodology to inspect if the reconstruction was indeed reaching super-resolution capabilities. We imaged commercially available calibrated polystyrene beads of various sizes, which are well-known to aggregate and form close-packed structures. Therefore, we can use the bead close contact distance as a proxy for the bead diameter. We measured the close contact distances of several beads for several sizes ranging from smaller than to several times the resolution limit, and also for two different objectives with different NAs. Although the method is somewhat subjective, we were careful to chose “spot centers” that had the smallest distances possible. Following this procedure, we noticed that maximum spot-to-spot center were indeed limited by the bead size, that is, in the 360 nm bead diameter we did not see 240 nm spatial fluctuations. The outcome of this procedure is shown Fig. S1 and the agreement between the nominal bead diameter and the retrieved diameter therefore confirms that the features observed in the blind-S^3^ reconstructions indeed correspond to physical features beyond the diffraction limit.

### III. SECTIONING ANALYSIS

We derive an analytic expression for explaining the depth sectioning ability of blind-S^3^. We assume two superposed Gaussian beams with electric fields *E* and 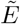 at two different wavelengths, and corresponding beam waist *ω*_0_ and 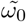, Rayleigh range *Z*_*R*_ and 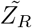. The two beams z-scans a continuous thin film containing Raman active centers *ρ* with a thickness *d*. The total signal detected *S*_*sect*_ can be calculated as:

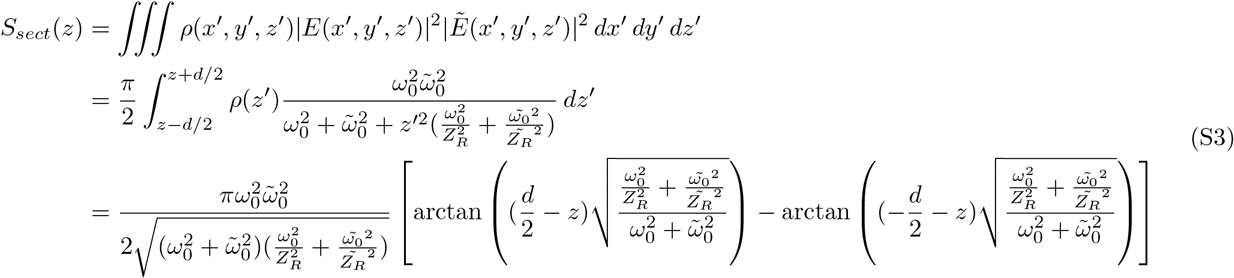

**FIG. S1.**
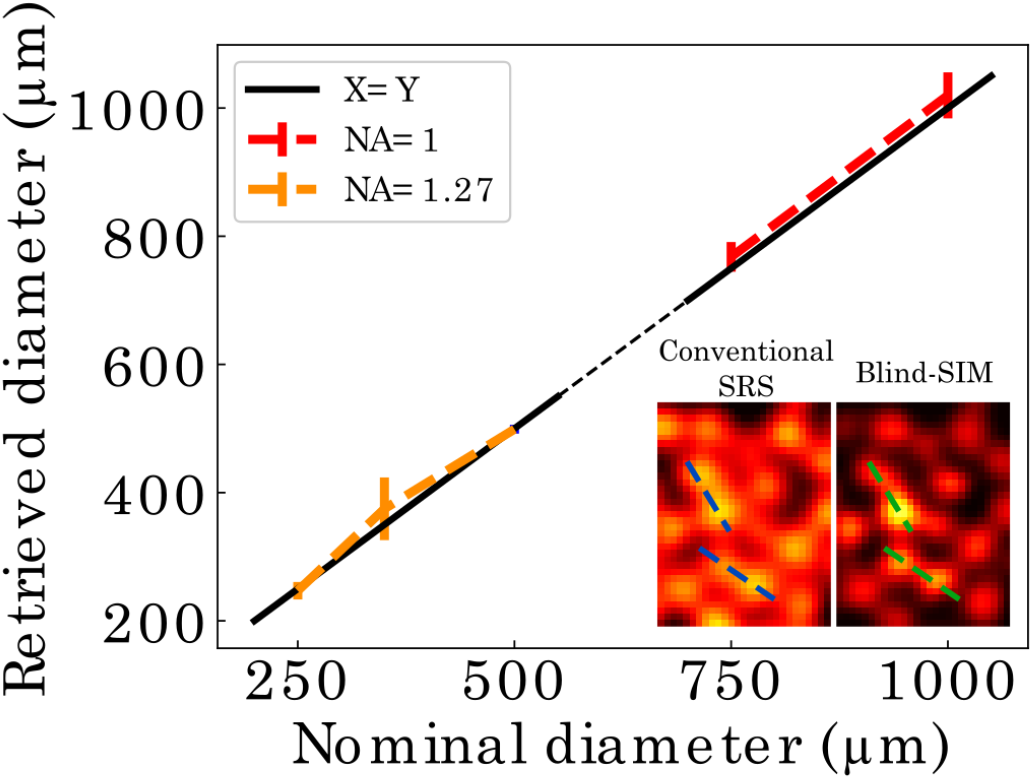
Transverse resolution analysis for blind-S^3^. Outcome analysis of the images of various close-contact beads pairs: We use close-contact distances as a proxy for the bead diameter. The inset shows representative images used for analysis, with the dashed lines representing some of the beads chosen for evaluation.

Comparison between the data and this simplified model shows a remarkable agreement, despite the fact that the rapid fluctuations of the speckle are not taken into account - that is, we do not use Eq. S2 for computing the average response, due to its high computational cost. In order to compare with the model, we have z-scanned a thin film in a blind-S^3^ acquisition procedure, and averaged over all realizations, for various FOV controlled by the number of independent macro pixels shown in the SLM (Fig. S2, left panel). We extract the full-width-at-half-maximum (FWHM) of each FOV and compare them with simulations using Eq. S3 Fig. S2 (middle panel): as seen in (Fig. S2, right panel), the agreement between the coarse approximation (which does not take into account the sharp fluctuations of the speckle, and neither the nonlinear polarization in the SRS process) and the data explain very well the dependence of the transverse FOV with the sectioning capabilities.

**FIG. S2.**
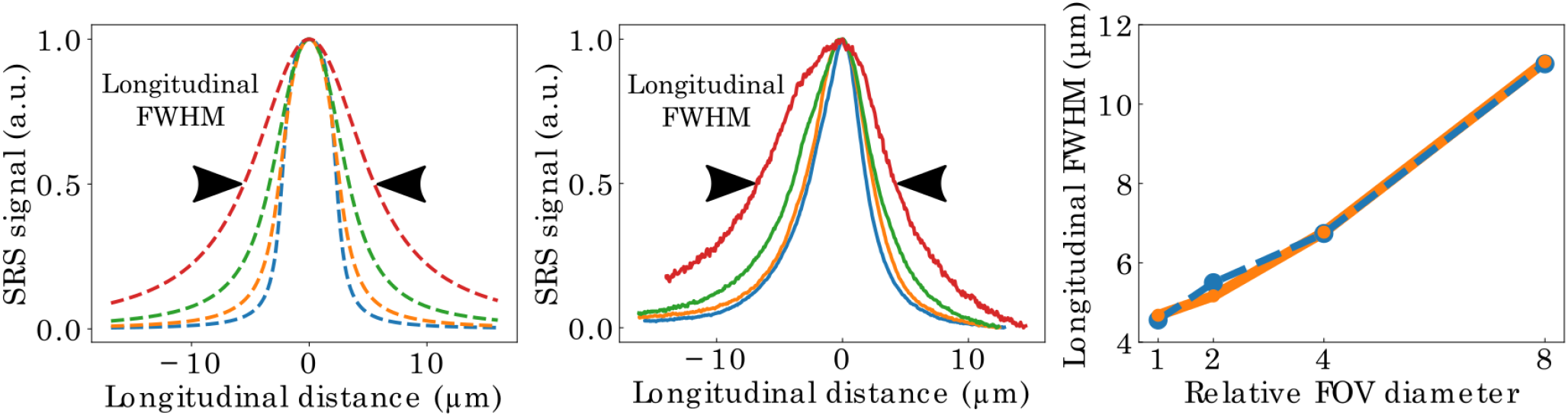
Depth sectioning analysis. (left panel) SRS signal of a z-scanned thin-film for various transverse FOV. Each curve is an average of multiple speckle realizations. (middle panel) Simulation of an SRS-equivalent process (see text and Eq. S3 development). (right panel) Extracted FWHM for the simulation (dash) and experimental data (line) vs the relative transverse FOV diameter.

### IV. LONGITUDINAL RESOLUTION

blind-S^3^ has high longitudinal resolution. We image a 2 layer structure of 500 nm polystyrene beads for multiple z-positions. The outcome of the SIM algorithm is presented in Fig. S3 for two layers separated by 1000 nm depth. The conventional SRS resolves the two layers, since they have different morphology, which is coherent with the resolution of conventional SRS 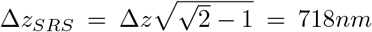 (with 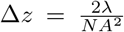. Therefore, we infer that the blind-S^3^ longitudinal resolution is similar to the conventional SRS given the similar morphology of the retrieved images.

**FIG. S3.**
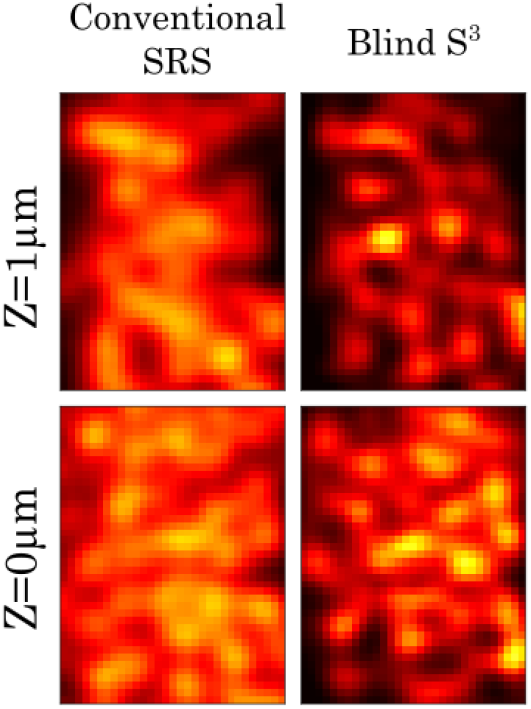
Longitudinal resolution results.

### V. OPTICAL LAYOUT

Figure S4 presents schematics of the optical layout.

**FIG. S4.**
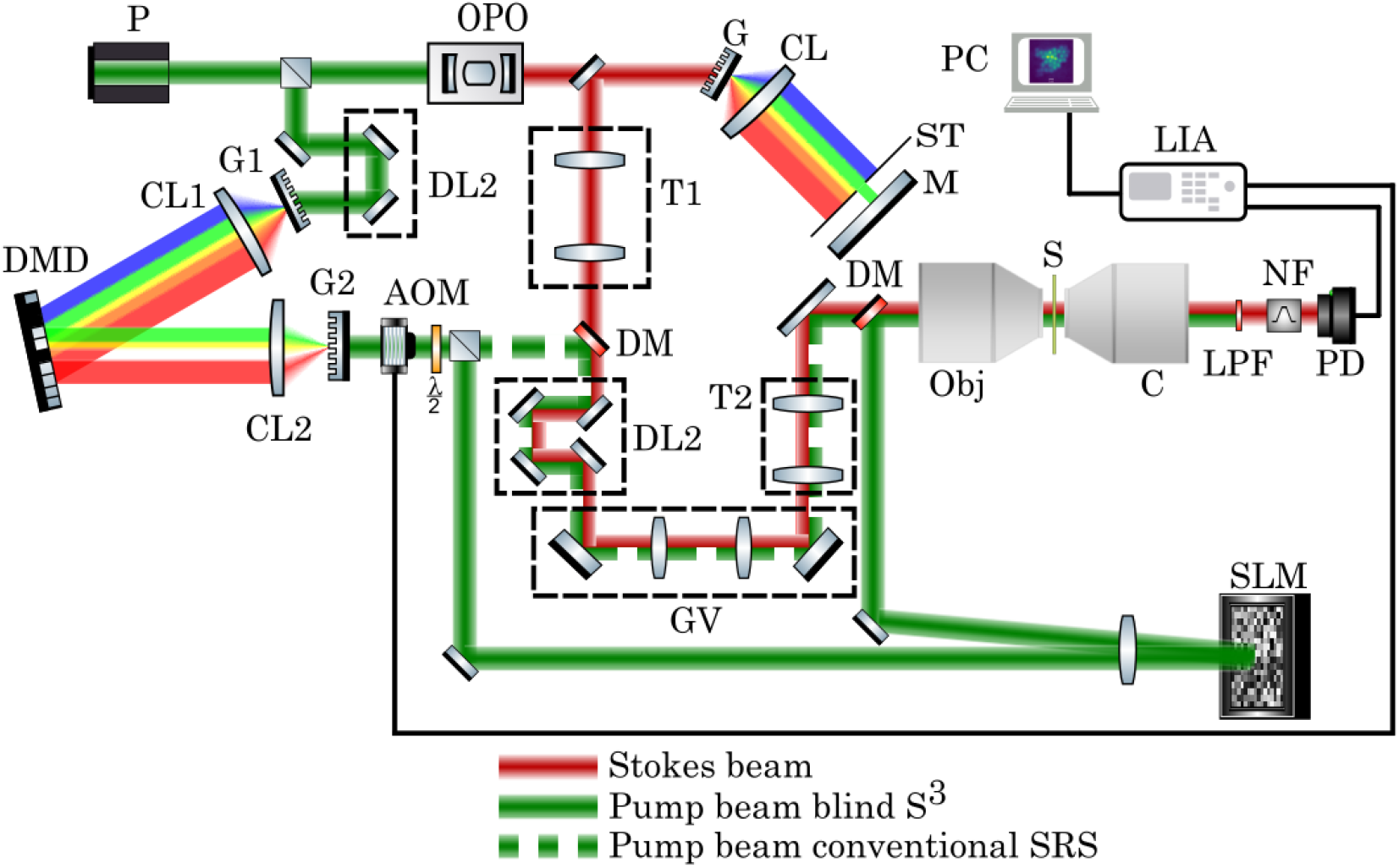
Detailed scheme of the setup used to perform experiments. Legend: P=Pump laser, OPO=optical parametric oscillator, DL= delay line, T=telescope, DMD=digital micromirror device, AOM=acousto-optic modulator, G=grating, CL=cylindrical lens, M=mirror, DM=dichroic mirror, SLM=spatial light modulator, Obj=microscope objective, C=condenseur, LPF=long-pass interference filter, NF=notch interference filter, PD=photodiode, LIA=lock-in amplifier, PC=personal computer.

## Notes

### Competing Interest Statement

The authors have declared no competing interest.

